# Integration of nuclear morphology and 3D imaging to profile cellular neighborhoods

**DOI:** 10.1101/2025.03.31.646356

**Authors:** André Forjaz, Donald Kramer, Yu Shen, Habin Bea, Margarita Tsapatsis, Jianglu Ping, Vasco Queiroga, Kyu Han San, Saurabh Joshi, Casey Grubel, Maria L. Beery, Irina Kusmartseva, Mark Atkinson, Ashley L. Kiemen, Denis Wirtz

**Affiliations:** Department of Chemical & Biomolecular Engineering, Johns Hopkins University, Baltimore, MD; The Johns Hopkins Institute for NanoBioTechnology, Johns Hopkins University, Baltimore, MD; Department of Pathology, The Sol Goldman Pancreatic Cancer Research Center, Johns Hopkins School of Medicine, Baltimore, MD; Department of Biomedical Engineering, Johns Hopkins University, Baltimore, MD; network for Pancreatic Organ Donors with Diabetes (nPOD) – confirm the abbreviation, Gainesville, FL; Department of Pathology, University of Florida, Gainesville, FL; Department of Functional Anatomy and Evolution, Johns Hopkins School of Medicine, Baltimore, MD

## Abstract

Nuclear morphology is an indicator of cellular function and disease states, as changes in nuclear size, shape, and texture often reflect underlying disease-related genetic, epigenetic, and microenvironmental alterations. For disease diagnosis, nuclear segmentation performed in 2D hematoxylin and eosin (H&E)-stained tissue sections has long represented the gold standard. However, recent advances in three-dimensional (3D) histology, which provide a more biologically accurate representation of the spatial heterogeneity of human microanatomy, has led to improved understandings of disease pathology. Yet challenges remain in the development of scalable and computationally efficient pipelines for extracting and interpreting nuclear features in 3D space. 2D histology neglects crucial spatial information, such as 3D connectivity, morphology, and rare events missed by sparser sampling. Here, through extension of the CODA platform, we integrate 3D imaging with nuclear segmentation to analyze nuclear morphological features in human tissue. Analysis of 3D tissue microenvironments uncovered critical changes in 3D morphometric heterogeneity. Additionally, it enables the spatial characterization of immune cell distribution in relation to tissue structures, such as variations in leukocyte density near pancreatic ducts and blood vessels of different sizes. This approach provides a more comprehensive understanding of tissue and nuclear structures, revealing spatial patterns and interactions that are critical for disease progression.

## INTRODUCTION

Digital pathology has revolutionized tissue analysis through high-resolution imaging and digitization of H&E-stained slides, enabling automated cellular and subcellular analysis ^1–11^. Traditional digital pathology focuses on 2D histological images, however such images are limited in their ability to capture the full complexity of tissue architecture and cellular composition, as spatial information is largely lost when observations are not extended from a single plane to three dimensions (3D) ^12–26^. A shift to 3D histology and analysis offers a more comprehensive view of the cellular microenvironment, revealing morphological features and pathological changes often missed in 2D studies ^13,21,22,27^.

Nuclear morphology closely mirrors cell phenotype, reflecting the functional state and identity of cells within their microenvironment and is a critical predictor of disease outcomes^28–30^. For example, heterogeneity in nuclear morphology within primary tumors can predict metastatic burden ^31–33^. However, the phenotype of a cell type is not static, it is dynamically influenced by the 3D microenvironment surrounding each cell, including mechanical forces, extracellular matrix composition, and interactions with neighboring cells ^34–38^. These factors collectively shape nuclear architecture, which in turn regulates gene expression, cellular function, and responses to pathological stimuli ^39,40^.

A critical research gap remains in developing robust pipelines to integrate nuclear morphology and tissue label features into 3D image datasets of large-volume samples, which is essential for understanding tissue organization and disease progression ^21,41–46^. This integration is crucial to analyze, for instance, immune hotspots within the 3D microenvironment, regions of high cellular activity and complex cell type organizations, which play pivotal roles in disease progression ^47–56^.

While intact 3D imaging techniques, such as light-sheet microscopy and tissue clearing, enable more direct 3D nuclear segmentation, these methods are limited in their ability to utilize large tissue volumes and susceptibility to morphological artifacts caused by suboptimal antibody penetration or low signal-to-noise ratios ^57–59^. In contrast, serial sectioning workflows support nuclear segmentation across extensive tissue volumes and preserve tissue blocks for subsequent profiling, enabling advanced immunostaining or multi-omics analyses to interrogate the tissue microenvironment, capabilities often lost during tissue clearing ^46,60,61^.

To overcome the limitations of 2D assessments of nuclear morphology, we introduce here a novel approach integrating 3D imaging with advanced nuclear and semantic segmentation techniques to analyze large tissue samples. Our methodology provides a unified pipeline for processing serial sectioning-derived 3D data and integrating cellular morphology with tissue-level annotations. 3D nuclear morphology mapping, together within the recently introduced CODA framework, provides high-resolution reconstructions of tissue architecture and enables the detection of alterations in morphology and cellular composition. By mapping nuclear morphology and cellular neighborhoods within the CODA framework, we identify structural alterations underlying pathologies and morphological attributes of disease progression, affording the ability to gain novel mechanistic insights into the spatial organization of pathological processes.

## RESULTS

### A novel workflow to integrate morphology and nuclear location in 3D space

A technical challenge in serial sectioning-based 3D tissue analysis is the precise alignment of nuclear segmentations to reconstruct spatial relationships among cells within their tissue microenvironment ^43,62,63^. Yet, while 2D nuclear detection and morphology analysis are well-established methods, analyzing nuclei in 2D sections alone misses critical spatial and contextual information as the cellular and non-cellular microenvironment of any given cell in a tissue is 3D. To address this challenge, we developed a pipeline for integration of nuclear segmentation coordinates and extensive nuclear morphology features with the CODA 3D reconstruction platform, allowing for comprehensive 3D assessments of tissue architecture and nuclear morphology ^45,60,64,65^.

To demonstrate this pipeline, we serially sectioned and 3D reconstructed a thick slab of human pancreatic tissue collected from a healthy organ donor and stained every fourth section with H&E (Fig. 1). First, we applied nonlinear image registration to H&E stained images to align and reconstruct serial images into a high-resolution digital 3D volume. Next, we used a semantic segmentation algorithm to label seven pancreatic tissue types identifiable in H&E at a resolution of 1 micron per pixel with 94.9% accuracy: acini, normal ductal epithelium, islets of Langerhans, vasculature, nerves, fat, and stroma. Finally, we performed nuclear segmentation through fine-tuning of the StarDist model for application to 20x-magnification whole slide images (WSIs), from its original optimization for 40x images (see supplemental methods for more technical information)^66^. The model achieved a measure of predictive performance F1 score of 0.89, 0.84, and 0.80 for three randomly chosen tiles of an independent testing set, demonstrating robust performance across diverse regions (Fig. S2a). Below, we will demonstrate that our pipeline is adaptable to any nuclear segmentation tool, such as StarDist or CellPose ^66–69^. Using these nuclear masks, we generated a list of 21 features describing nuclear shape, size, stain intensity and heterogeneity (Table S1), which are automatically generated for nuclei detected with our pipeline.^70^ Additionally, we extended the use of CODA to enable registration of nuclear coordinate features into the 3D volume for integrated cellular and tissue-level correlation (Fig. 2b, top and bottom).

**Fig. 1.**
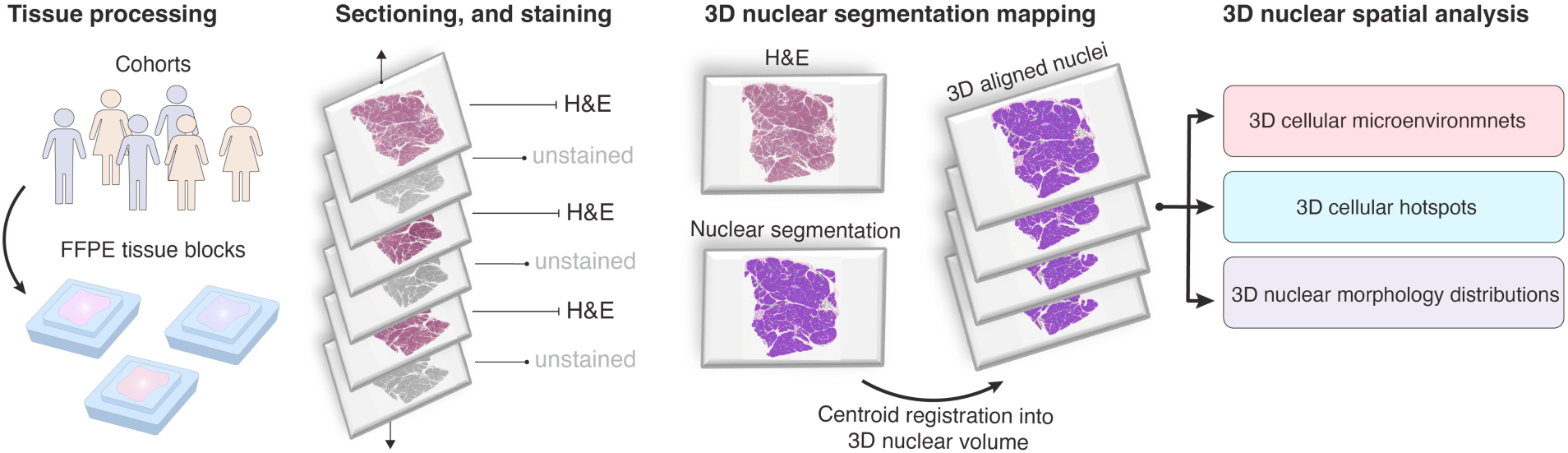
Construction of 3D nuclear segmented and microanatomically labelled tissue cohorts. A FFPE tissue is resected and serially sectioned. Every fourth section was stained H&E and digitized at 20x resolution. Tissue- and nuclear-level segmentation is performed on the digitized H&E images and registered into spatially aligned 3D tissue volumes. CODA-deep learning model is applied to automatically label nuclei according to their anatomical labels in 3D. Comprehensive 3D nuclear spatial analyses are conducted to profile the tissue architecture, including evaluation of 3D cellular microenvironment, identification of 3D cellular hotspots, and characterization of nuclear morphology distributions in tissue volumes.

**Fig. 2.**
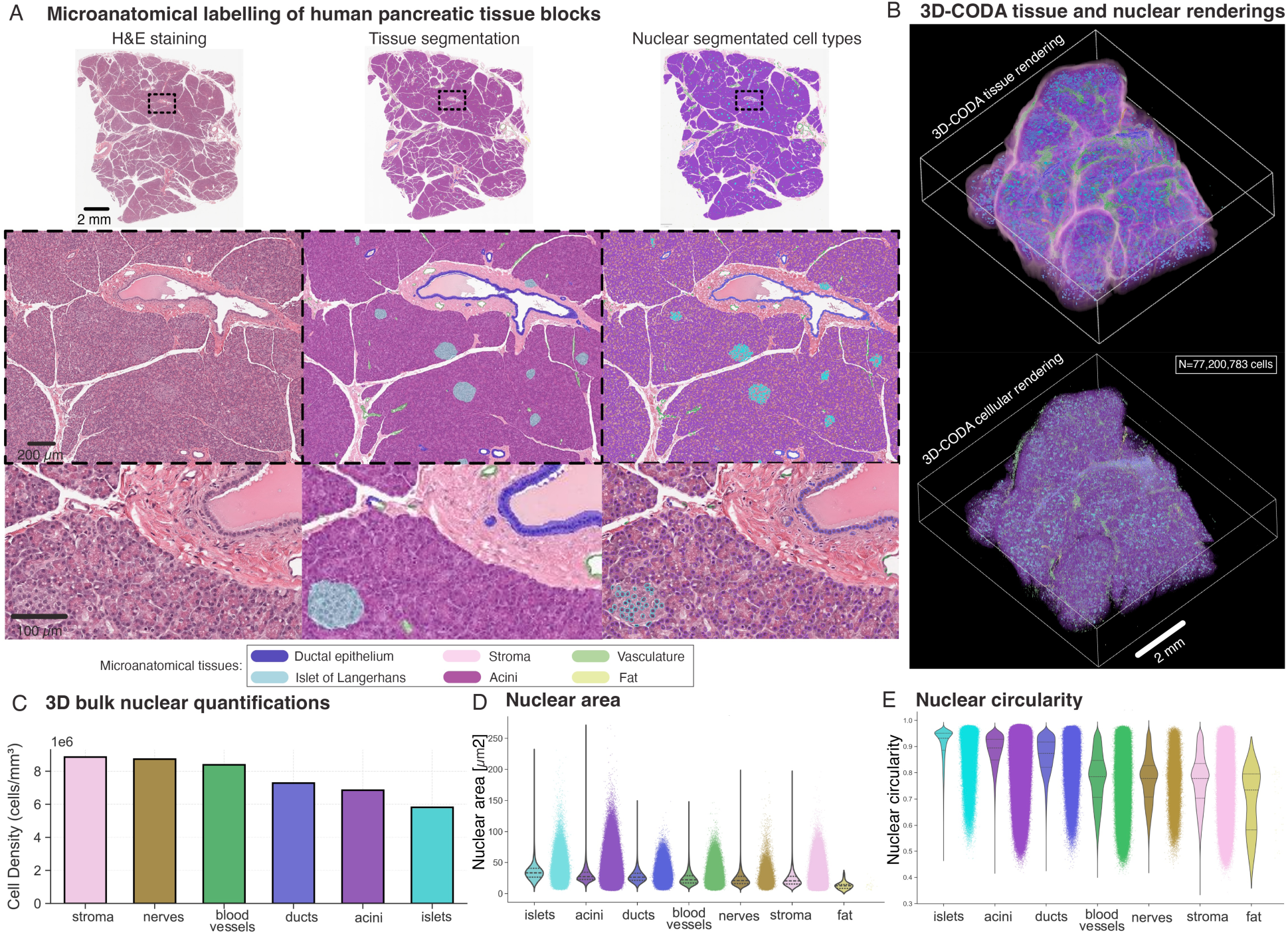
Reconstruction of nuclear morphology in three-dimensional tissues. (**a**) CODA semantic segmentation was used to annotate seven microanatomical components of H&E-stained pancreatic tissue sections: including ductal epithelium, acini, islets of Langerhans, blood vessels, nerves, stroma, fat, and background. Nuclear segmentation coordinates were integrated with CODA microanatomical labels, enabling precise classification of individual nuclei within their tissue context. (**b**) 3D-CODA platform reconstructed the spatial architecture of tissue components (top panel), and generated the 3D rendering of 77 million nuclear centroids, each color-coded according to its CODA microanatomical label (bottom panel). (**c**) Quantitative assessment of nuclear features, including cell density (# nuclei/volume) and nuclear morphology (size and shape) of cells, both stratified by microanatomical labels of the pancreas. Cell density across different tissues of the pancreas sample. (**d**) Nuclear areas stratified by microanatomical labels; we show both violin plots and distributions for each tissue type). (**e**) Bulk nuclear circularities analysis, enabling comparative analysis of nuclear morphology across tissue types.

By integrating nuclear segmentation with 3D CODA-based tissue segmentation and coordinate registration, we generated high-resolution 3D maps of nuclear morphologies by cell type (Fig 2A)^60^. This enabled tissue reconstruction of the human pancreatic tissue (Fig. 2b, top) and nuclear segmentation reconstruction in that tissue (Fig. 2b, bottom), preserving spatial relationships between nuclei and tissue structures such as ducts, nerves and blood vessels, and enabling the analysis of nuclear clustering, and local cell density by cell type. Below, we demonstrate how this integrated tissue and cellular data enables diverse analyses, including classification of cell types using 3D neighborhood information and correlation between 3D nuclear information and local microenvironment in large-volume tissues.

### 3D nuclear segmentation reveals diverse tissue and cellular heterogeneity

This initial reconstructed sample measured approximately 16 mm × 14 mm × 2.02 mm (volume: 452.5 mm^3^) and contained approximately 77 million nuclei, 100x-fold more cells than are present in a single WSI of the 3D dataset. Bulk cellular and volumetric quantifications revealed significant heterogeneity in tissue composition (Fig. S2b). Structures such as stroma and nerves showed notably lower cell density (fewer cells per unit volume), while acini and islets of Langerhans were markedly denser (Fig. 2c, left).

Nuclear morphological parameters, including area and circularity, were extracted for each of the 77 million cells and revealed substantial variability across tissue types (Fig. 2c, middle, left; Fig. S2d). In total, 21 nuclear features were quantified, and correlation analyses underscored the broad range of nuclear characteristics at single-cell resolution (Fig. S2c). Among these tissues, cells in islets of Langerhans displayed a distinct morphological profile, exhibiting high circularity (mean = 0.90, SD = 0.06, CV = 0.07) and relatively uniform shapes, yet a considerable degree of variation in nuclear area was noted (SD = 12.2 µm², CV = 0.35). Despite this variability, islets also exhibited the largest mean nuclear area (34.3 µm²). By contrast, acinar cells demonstrated a more homogeneous size distribution (SD = 9.9 µm², CV = 0.34), with high circularity (mean = 0.88, SD = 0.06, CV = 0.07) and an intermediate mean area (29.1 µm²), suggesting a comparatively consistent architectural organization. Cells in blood vessels, nerves, pancreatic ducts, and the stromal compartment presented intermediate levels of heterogeneity in both shape and size, indicating moderate structural complexity. Fat cells exhibited notable morphological variation, although their limited sample size prevented definitive conclusions about their overall distribution. These results demonstrated how our novel workflow can analyze exceedingly large numbers of nuclei in the same sample and associated complex landscape of 3D nuclear morphological diversity of the human pancreas.

### Identification of leukocytes based on nuclear morphology and 3D associated microenvironment

To demonstrate the utility of determining nuclear morphology in 3D tissues, we developed an additional workflow to determine whether we could classify cell types using nuclear morphology features alone, demonstrated here through identification of CD45+ leukocytes. A subset of 249 leukocytes and 249 non-leukocytes were manually annotated using QuPath and matched to corresponding nuclear segmentation features^71^. A random forest classifier was trained using hyperparameter tuning, with 5-fold cross-validation optimized key parameters (number of trees, maximum depth, minimum leaf size) using MATLAB.^72^ The best performing hyperparameters were identified and used to train the final model.^73–75^ This model achieved an accuracy of 89.04% on independent testing of images (Fig. S3a). The classifier trained on 10 sections was then applied to the entire 3D volume (101 sections), allowing for the generation of 3D leukocytes maps in the human pancreas sample (Fig. 3a).

**Fig. 3.**
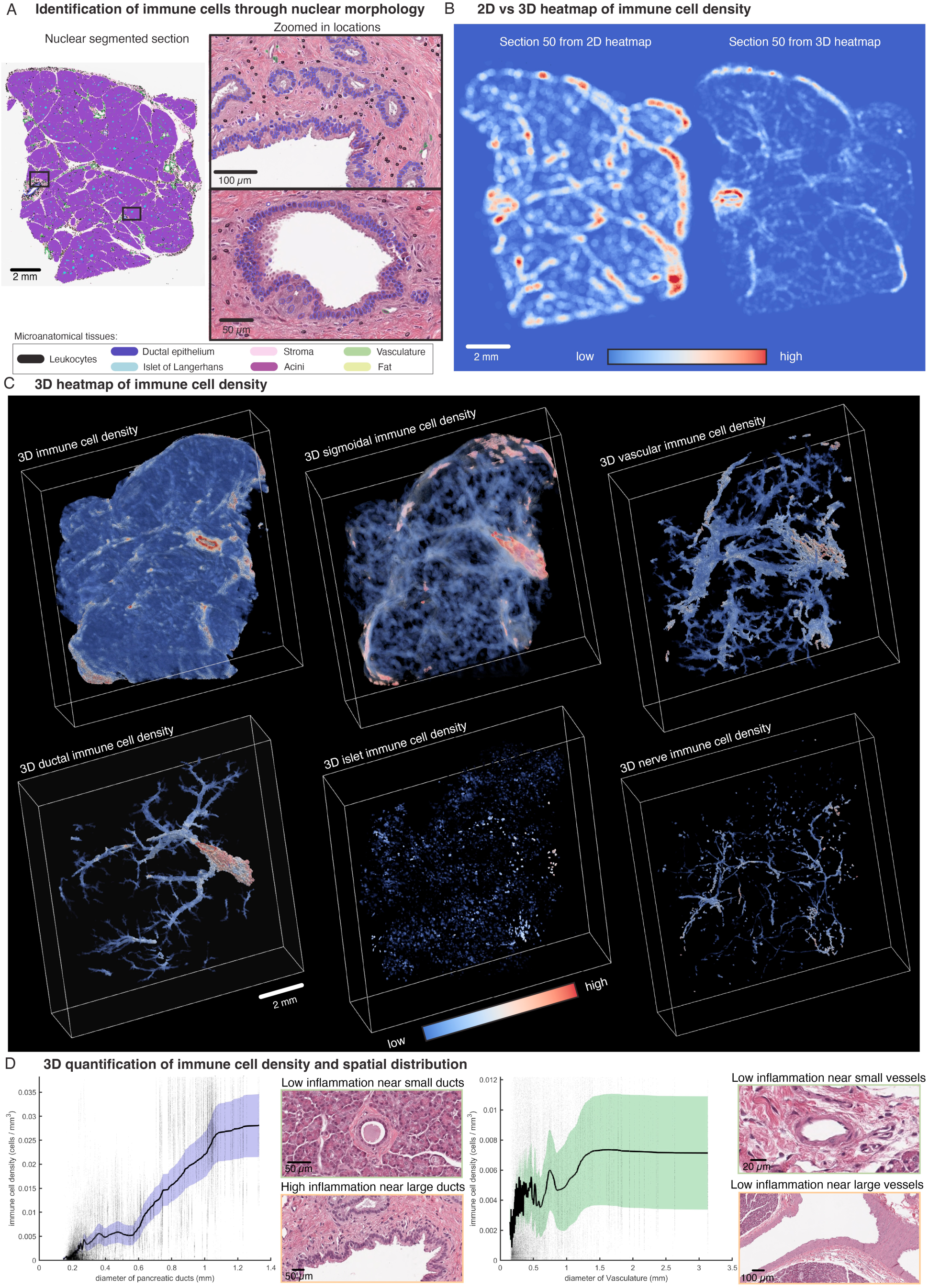
Three-dimensional quantification of leukocyte cell densities and their spatial interaction with surrounding cellular components in human pancreatic tissue. **(a)** Leukocytes were identified by manually selecting a representative subset of leukocytes and training a random forest classifier to predict the remaining leukocyte population across the 3D dataset. **(b)** Leukocyte heatmap of a single 2D section revealed multiple leukocyte hotspots, whereas the corresponding 3D leukocyte heatmap of the same section revealed more spatially coherent leukocyte hotspots, providing a more comprehensive view of hotspot distribution in the entire 3D pancreatic sample. **(c)** 3D heatmap visualization depicting leukocytes proximity to distinct tissue components, enabling the assessment of spatial interactions. **(d)** Quantitative analysis of leukocyte density in the vicinity of the ducts showed an increase in leukocyte density around larger pancreatic ducts. Example of a small duct with low inflammation (left, top panel), and a large duct with high inflammation (left, bottom panel). Similarly, leukocyte density was moderately elevated near larger blood vessels compared to smaller vessels. Example of a small vessel with low inflammation (right, top panel), and a large vessel with higher inflammation (right, bottom panel).

We hypothesized that 3D analysis would reveal the true 3D topology of leukocyte hotspots in the human pancreas compared to 2D traditional assessments, as 2D methods are prone to distortions from sparse sampling, which can misrepresent spatial density and connectivity. Comparing a randomly selected 2D single-section immune hotspots (Fig. 3b, left) with 3D volume immune hotspots (Fig. 3b, right) confirmed this hypothesis, showing more localized and precise patterns in 3D. 3D leukocyte cell density heatmaps identified hotspots near stromal, vascular, and epithelial regions, while cold spots were predominantly observed in islets and nerves (Fig. 3c) ^45^. These spatial patterns differed significantly from 2D analyses, which often over- or under-represented leukocyte densities due to sampling bias.

We further exploited our 3D dataset and found through quantitative analysis that a significantly higher leukocyte density was present around larger pancreatic ducts compared to smaller ducts, indicating a size-dependent inflammatory response (Fig. 3d, left). Similarly, leukocyte density was moderately elevated near larger vessels relative to smaller ones, suggesting a link between vessel size and inflammatory cell recruitment (Fig. 3d, right). Such assessments are not possible using traditional 2D tissue sampling techniques.

### The three-dimensional cellular and non-cellular microenvironment around each cell of a tissue

Our integration of 3D CODA mapping with nuclear segmentation opens the opportunity to assess the cellular and non-cellular composition of the microenvironment in the vicinity of each cell in the entire 3D tissue sample. To accurately assess the true (unbiased) composition of the microenvironment surrounding each cell, 3D analysis is essential. 2D methods fail to capture the full spatial context, as they are limited by sparse sampling and the inability to account for out-of-plane interactions, leading to incomplete or misleading representations of cellular neighborhoods.

To determine the 3D microenvironment for each cell, we first generated spheres of radii between 16 and 128 µm around each individual nucleus in the pancreatic sample (Fig. 4a). Then, we generated heatmaps of the 3D microenvironment composition (Fig. 4b), which revealed a decline in content for sparser tissue components, such as islets, leukocytes, and nerves, and an increase in more abundant tissues, such as acini, as the microenvironment radius increased. This was consistent with qualitative observations in histological slides (Fig. 4c). Larger cell populations became more predominant with increasing microenvironment size, a trend observed across all cell types (Fig. 4d).

**Fig. 4.**
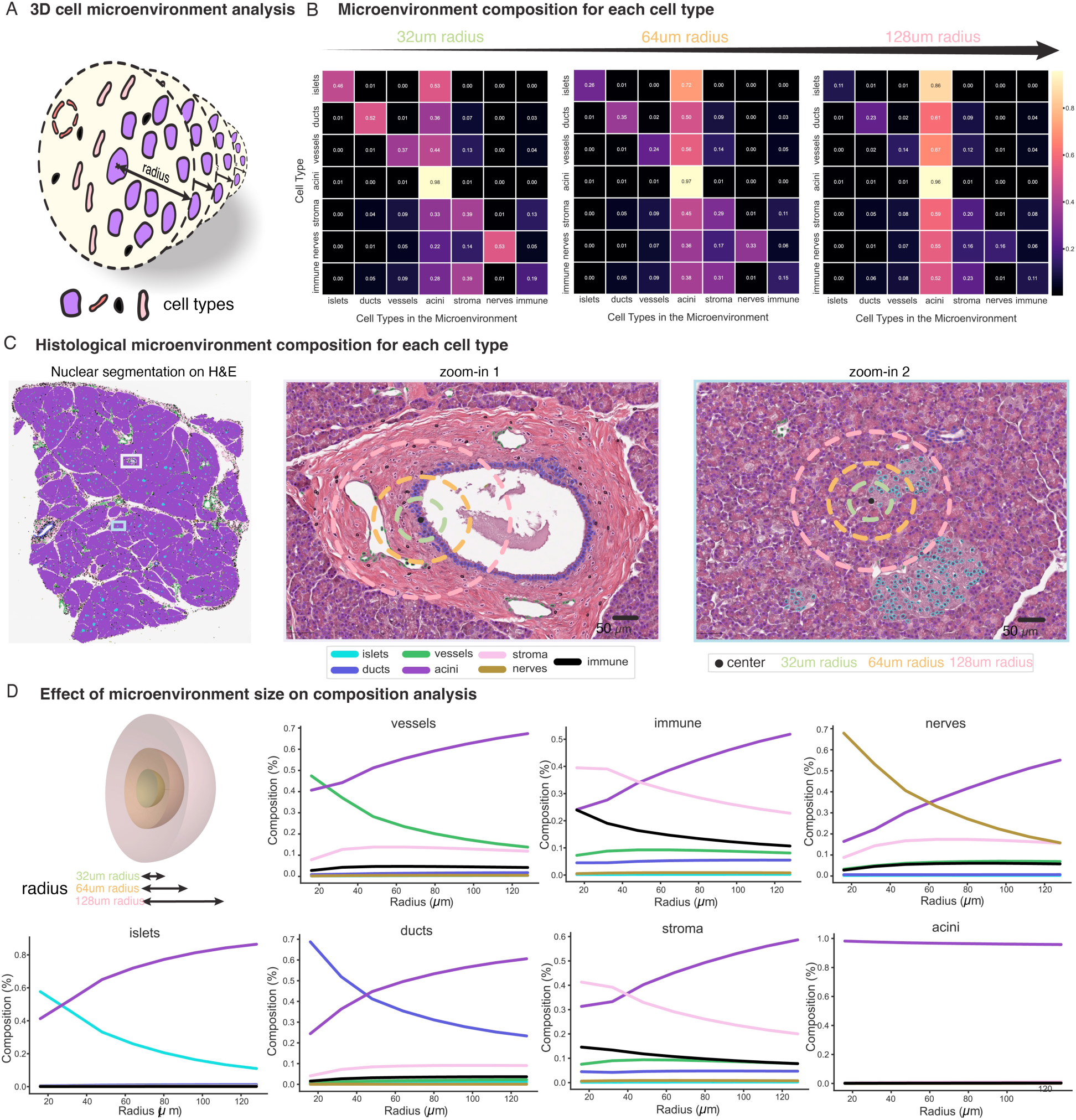
Three-dimensional assessments of the cellular microenvironment for each individual cell of a human pancreatic tissue. (**a**) For each cell within the 3D tissue volume, spherical regions were generated at defined radii to quantify the number and types of neighboring cells, allowing localized microenvironment profiling. (**b**) Quantitative assessment of the cell type composition within these spherical microenvironments was performed across varying radii. Smaller radii sizes were able to capture smaller and localized cell populations, whereas larger radii were more prone to capturing more abundant cell populations in their vicinity, revealing differences in microenvironmental complexity based on spatial scale. (**c**) Visualization of different radius sizes overlaid in H&E-stained histological slides, alongside with nuclear segmented and anatomically labelled cells, illustrating how the chosen radius impacts microenvironmental analysis. (**d**) Tissue composition analyses of the cellular microenvironment for each cell type across increasing radii. For each cell type, the composition of the surrounding cell was measured as the radius increased, highlighting shifts in neighboring cell populations as the radius expands. Since the composition of the healthy human pancreas is dominated by acini, the composition of the cellular microenvironment around each type of cell is eventually dominated by acini for large radii around the cell.

## DISCUSSION

This study integrates 3D imaging at single-cell resolution with nuclear segmentation within the CODA 3D cellular imaging platform. As a testbed for this integration, we used the human pancreas, one of the most complexes organs ^46,76,77^. While traditional 2D analyses have historically been the gold standard for solid tissue histology and provide valuable insights, they are intrinsically limited and do not capture the full spatial complexity of the tissue microenvironment. Our 3D approach overcomes this limitation by enabling the extraction of nuclear morphology, alongside spatial characterization of leukocyte distribution within tissues. Notably, we observed increased leukocyte density in proximity to larger pancreatic ducts and blood vessels, a pattern that would be difficult to resolve using 2D histology.

By integrating nuclear segmentation with semantic classification, this method enables the identification of 3D cellular architecture and spatial density gradients (i.e., heatmaps), which are often overlooked in conventional 2D analyses. This framework also facilitates spatially resolved characterization of tissue heterogeneity and cellular interactions at single-cell resolution, providing a more comprehensive understanding of microenvironmental organization. Such capabilities are particularly relevant for studying cancer pathogenesis, where the interplay between spatial organization, immune infiltration, and tumor progression is critical.

Despite these advances, challenges remain. The computational demands of 3D segmentation and analysis are significant, requiring robust hardware and efficient algorithms. Additionally, reliance on H&E-stained images limits the granularity of cellular feature extraction. Integrating multi-omics data could enhance the precision and biological relevance of our analyses^64^. Future studies should validate this approach across diverse tissues and disease models. Expanding this framework to study dynamic processes, such as tumor evolution or immune responses, could further advance our understanding of disease progression.

In conclusion, our 3D nuclear segmentation approach provides a powerful framework for analyzing large tissue complexity, offering insights into disease mechanisms. Future work should focus on broader applications, multi-modal data integration, and overcoming current limitations to solidify 3D imaging as a gold standard of digital pathology.

## Supplemental figures and captions

**Fig. S2.**
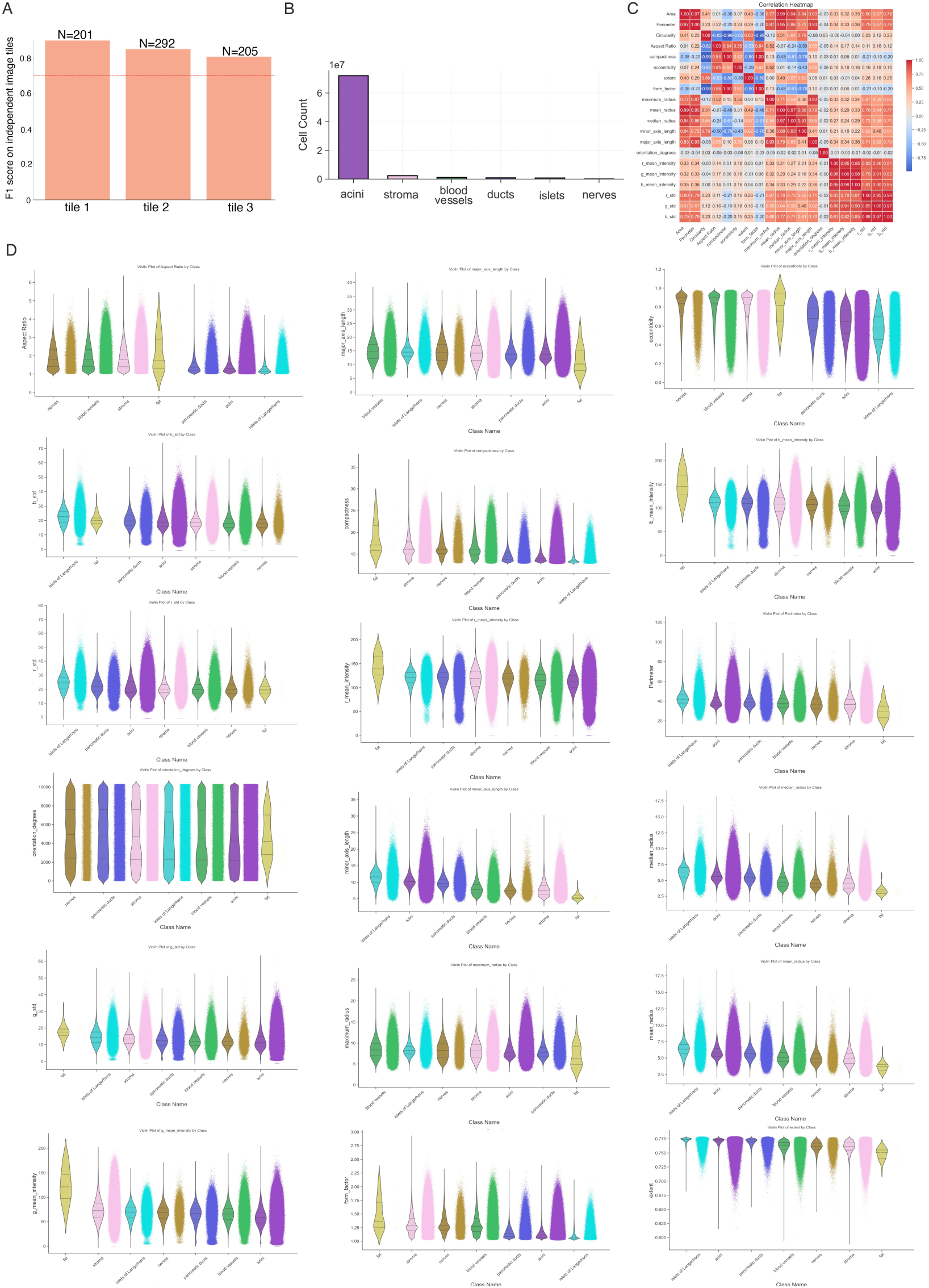
Quantification and validation of nuclear morphology. (**a**) Subset of tiles was used to validate the accuracy of the nuclear segmentation, yielding an F1 score of 0.89, 0.84, and 0.80 (**b**) Segmented nuclei were integrated with the 3D-CODA platform, with each color-coded according to its CODA microanatomical labels. Cell counts for each annotated cell type were extracted. (**c**) Correlation plot of the nuclear features highlights the degree of variation across different nuclear attributes. (**d**) Nuclear morphology feature distributions were analyzed in relation to CODA tissue labels, revealing distinct variations in nuclear properties across different tissue compartments.

**Fig. S3.**
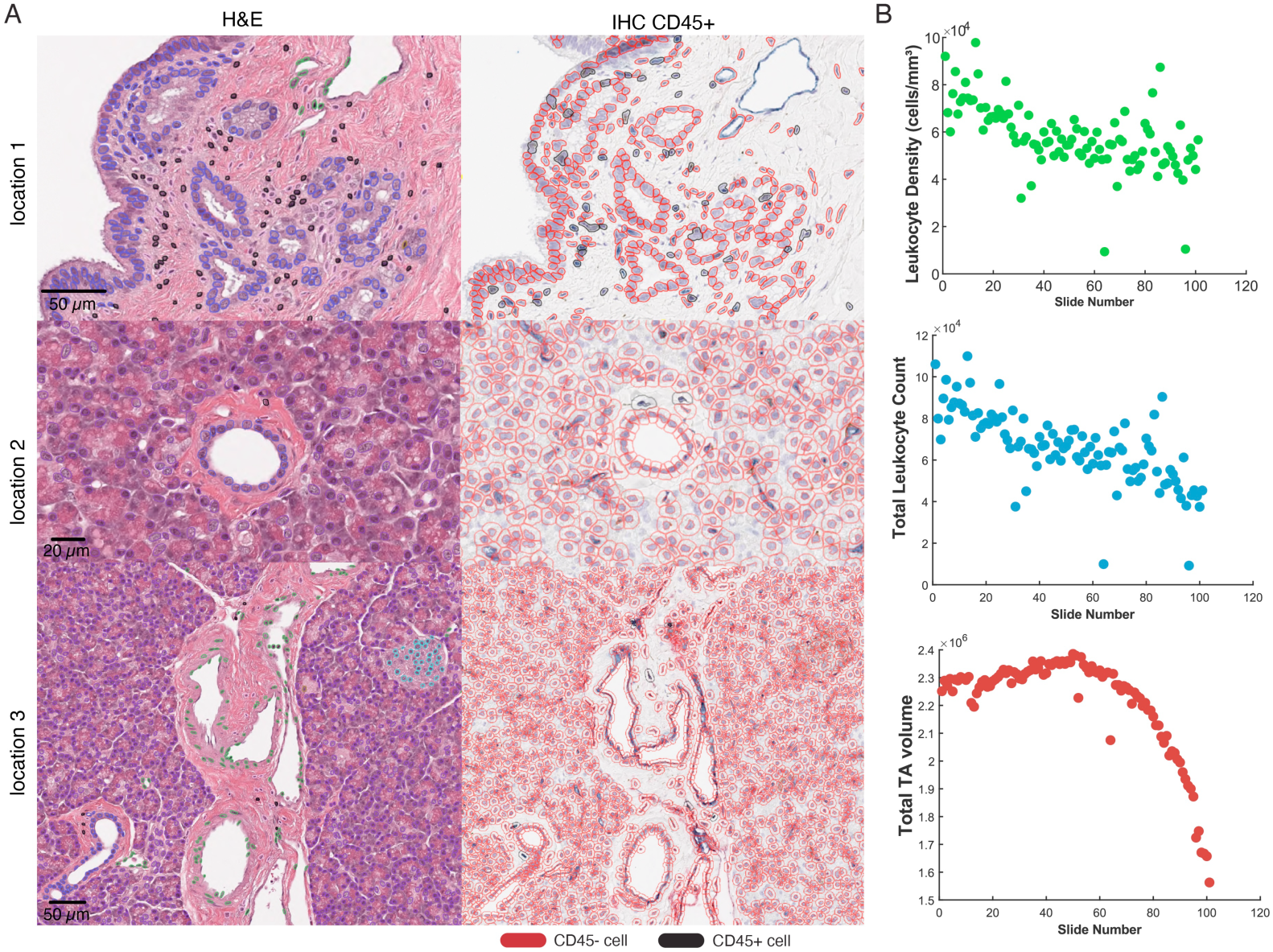
(**a**) Quantitative analysis of total leukocyte cell counts across individual tissue slides within the 3D tissue block showed an accuracy of 89.04%. (**b**,top) Total leukocyte cell density across the individual tissue slides within the 3D tissue block. (**b**,middle) Total leukocyte cell counts stratified by tissue slide. (**b**,bottom) Total tissue area (TA) across individual tissue slides within the 3D tissue block.

**Table S1.** Nuclear features extracted from H&E images. For each segmented nucleus, 21 morphological and intensity-based features were computed to profile cellular heterogeneity. A subset of nuclei was annotated to train a classifier for leukocyte identification based on these features.

| Feature Name | Description |
| --- | --- |
| Centroid x | X-coordinate of the nucleus centroid. |
| Centroid y | Y-coordinate of the nucleus centroid. |
| Area | Area of the nucleus (in pixels or $\mu\text{m}^2$, depending on resolution). |
| Perimeter | Perimeter of the nucleus. |
| Circularity | Measure of how circular the nucleus is (closer to 1 = more circular). |
| Aspect Ratio | Ratio of the major axis length to the minor axis length. |
| Compactness | Measure of how compact the nucleus is. |
| Eccentricity | Measure of how elongated the nucleus is (0 = circle, 1 = line). |
| Extent | Ratio of the nucleus area to the bounding box area. |
| Form factor | Shape descriptor based on area and perimeter. |
| Maximum radius | Maximum distance from the centroid to the nucleus boundary. |
| Mean radius | Mean distance from the centroid to the nucleus boundary. |
| Median radius | Median distance from the centroid to the nucleus boundary. |
| Minor axis length | Length of the minor axis of the nucleus. |
| Major axis length | Length of the major axis of the nucleus. |
| Orientation degrees | Orientation of the nucleus (in degrees). |
| R mean intensity | Mean intensity of the nucleus in the red channel. |
| G mean intensity | Mean intensity of the nucleus in the green channel. |
| B mean intensity | Mean intensity of the nucleus in the blue channel. |
| R intensity std | Standard deviation of intensity in the red channel. |
| G intensity std | Standard deviation of intensity in the green channel. |
| B intensity std | Standard deviation of intensity in the blue channel. |

## MATERIALS AND METHODS

### Tissue acquisition, processing and imaging

Pancreatic tissue was collected from an organ donor through the network for Pancreatic Organ Donors with Diabetes (nPOD) in accordance with existing federal and state regulations and with approval from University of Florida IRB. The formalin-fixed, paraffin-embedded (FFPE) pancreatic tissue block was serially sectioned at a thickness of 4 µm throughout the entire block, producing 404 tissue sections. Every fourth sections were H&E stained and scanned at 20x resolution (∼0.5 micron/pixel) using a Aperio SC2 slide scanner (Leica Biosystems Imaging, Inc). NDPI and scanned whole slide image SVS files were converted to tiff images (1 micron/pixel) for 3D-CODA tissue segmentation using the openslide ^78^.

### Software and Hardware Setup

The computational workflow was implemented using the CODA, combined with Python 3.9 and MATLAB ^60,72^. Open-source tools such as StarDist for H&E nuclear segmentation and QuPath for annotation and validation ^66,71^. Workstations equipped with an NVIDIA RTX 4090 GPU and 128 GB of RAM were used for computational processing. MATLAB was used for alignment of images and nuclear centroids into 3D volumes ^45^.

### CODA microanatomical tissue-level labelling

CODA semantic segmentation was used to identify cell types in H&E, including blood vessels, nerves, vasculature, acini, epithelium ducts, islets of Langerhans, among other in all WSIs ^60,79^. The model was trained using annotations from 5 images, and tested on an independent annotated image, achieving an overall accuracy of 94.9%.

### Alignment of 2D serial histology into 3D labelled tissue maps

CODA nonlinear image registration was used to align the serial histology and enable 3D reconstruction of microanatomical structures. Registration was calculated at a resolution of 8 micron per pixel and applied to the higher resolution (1 micron per pixel) segmented images to generate 3D labelled image stacks.^60^

### Nuclear segmentation

Nuclear segmentations was performed on 101 H&E stained pancreatic tissue images, using the Stardist pipeline ^66^. A pre-trained StarDist model, originally trained on 40x resolution H&E images, was fine-tuned for 20x resolution NDPI and SVS image files. To finetune the model, we annotated 25 tiles with 256×256 dimensions for training, followed by subsequent testing on 3 independent tiles. Finetuning of the pretrained model was optimized through adjustment of hyperparameters such as the learning rate, training epochs, and data augmentation. Tiles were selected from diverse regions of the sample to ensure morphological heterogeneity. Fine-tuned model performance was validated using precision, recall, and F1 score. For segmentation, WSIs were divided into 4096 x 4096 pixel tiles with an overlap parameter of 128 pixels to minimize edge artifacts. Segmentation results, including nuclear contours and centroids, were saved as JSON files for downstream analysis.

### Feature extraction from nuclear contours

From the nuclear centroids and contours of each segmented nucleus, a wide range of morphological and intensity-based features were extracted to characterize cellular and nuclear properties (Table S1). These features provide quantitative insights into nuclear shape, size, and staining patterns, which are critical for understanding cellular behavior and tissue organization. Here, we extracted cellular parameters such as area; long and short axis length; aspect ratio; eccentricity; diameter; average red, green, and blue intensities; standard deviation of red, green, and blue intensities; cell ID, slide number, and xyz coordinate.^80^

### Registration of 2D nuclear segmentation into 3D space

To reconstruct the 2D nuclear segmentation masks into 3D space, the CODA registration integration algorithm was extended to handle nuclear segmentation format data. Following CODA registration of the serial histology, the registration transforms were used to register the xy coordinates output following 2D Stardist segmentation into 3D tissue space. This step enabled integration of nuclear coordinate data with the tissue-type segmentation output, enabling identification of major cell types in 3D space.^45^ Tissue labeling data were integrated to ensure anatomical consistency, preserving spatial relationships between nuclei and tissue structures.

Once registered, the 2D nuclear segmentations were concatenated into a 3D volume, with each nucleus assigned to a unique cell ID. This ID linked each nucleus to its spatial coordinates and previously extracted morphological features, such as area, eccentricity, and intensity metrics.

### Measurement of bulk cellular and volumetric quantifications

Using the generated 3D tissue and cellular volumes, bulk quantifications were extracted to characterize tissue composition and cellular distribution. Volumetric data for each tissue component was calculated by summing the voxels corresponding to each label and adjusting for voxel size. Bulk cellular information of each microanatomical label can be extracted by integrating the 3D tissue-labelled volume with the spatial locations of cellular labels in the 3D cellular volume, enabling in silico analysis of microanatomical regions.

### Distinguishing Cell Types Using Nuclear Morphology Features

We demonstrated the ability of this algorithm to predict cell types using nuclear morphology through identification of CD45+ lymphocytes. A subset of lymphocytes was manually annotated in QuPath by researchers with expertise in histology. Annotated cell coordinates were exported as GeoJSON files, capturing their spatial coordinates and contours. The GeoJSON data was converted to a JSON file. The annotated cells were then processed to extract nuclear morphology features (e.g., area, eccentricity, intensity metrics) and stored in a pickle file. Using the morphology features of the annotated immune cells as a reference, a random forest classifier was trained to extrapolate immune cell identities across the entire 3D tissue block. The classifier was applied to all nuclei in the dataset, and the predicted immune cells were mapped onto the 3D volume. This enabled the generation of 3D renderings of immune cell infiltration, identifying regions of high immune activity (hot spots) and low immune activity (cold spots) within the tissue microenvironment. Separate images were CD45 Immunochemistry (IHC) stained to validate the performance of the classifier.

### Assessment of 3D nuclear cell morphology microenvironment

To evaluate the microenvironment of each cell nucleus, spherical volumes with different radii were generated for all nuclei in the dataset. Cellular, volumetric, and nuclear morphological features within each sphere were computed and tabulated according to microanatomical labels. Each row in the resulting tables represented a unique cell ID, with columns detailing cell counts, volumetric data, and nuclear morphology metrics for each label.

### 3D heatmap generation for visualizing intra- and inter-variability in tissue blocks

To visualize variability in cellular content, spherical volumes of 144 micron radius were generated for each voxel in the biospecimen. The number of nuclei within each sphere was summed, creating a 3D matrix of local nuclear densities. This matrix was used to generate color-graded heatmaps, overlaid on the 3D tissue volume, to highlight regions of high and low cellular density.

### Statistical considerations

Metrics within and between cohorts were compared using median, mean, standard deviation, and interquartile range. CODA segmentation model accuracy was determined through annotation of an independent testing image and calculation of per-class precision and recall. Nuclear segmentation model accuracy was determined through calculation of F1 score. No additional statistical analyses were performed in this study.

## Code and data availability statement

The data analyzed here is available from the corresponding author upon request. The code used to generate the 3D tissue maps is available on the following GitHub: https://github.com/ashleylk/CODA. The code used to generate the 3D nuclear morphology maps will be available upon publication.

## Author contributions

A.F., A.L.K. and D.W. conceived the project. M.A. C.G., M.L.B, I. K. collected and processed the human pancreas samples. A.F., D.K., Y.S., A.L.K, V.Q, R.T., L.P., H.B., S.J., and K.H.S. conducted image analysis and heterogeneity quantifications. A.F., A.L.K., and D.W. wrote the first draft of the manuscript, which all authors edited and approved.

